# Scanless temporal focusing enables high-speed three-dimensional quantitative phase microscopy

**DOI:** 10.64898/2026.03.04.709629

**Authors:** Yuechuan Lin, Xiang Zhang, Rebecca E Zubajlo, Zahid Yaqoob, Brian W. Anthony, Peter T. C. So

**Author notes:** Equal contribution (Xiang Zhang, Rebecca E Zubajlo). Current address: McGovern Institute for Brain Research, Massachusetts Institute of Technology, Cambridge, MA, 02139, USA. Corresponding to: Peter So, Zahid Yaqoob, Brian W. Anthony.

## Abstract

Quantitative phase microscopy (QPM) enables label-free imaging of structure and dynamics in biological and physical systems, yet achieving high-speed three-dimensional (3D) QPM with strong optical sectioning remains a central challenge. Here, we introduce a single-shot reflection-mode temporal focusing QPM (TF-QPM) that provides sub-micron optical sectioning without needs of any mechanical scanning or multiplexed acquisitions. By extending temporal focusing beyond its conventional use in multiphoton fluorescence microscopy, TF-QPM enables diffraction-limited label-free phase-sensitive volumetric imaging with 402 nm lateral and 920 nm axial resolution, markedly reduced speckle noise, and depth-resolved imaging at 3,709 Hz frame rate —an order of magnitude faster than most existing techniques and currently only limited by the camera speed. The resulting spatiotemporal phase sensitivity enables precise 3D tracking of particle motion and quantitative characterization of fast dynamics in complex and anisotropic media. For tissue imaging applications, TF-QPM achieves histology-level resolution in intact samples and supports pixel-level virtual staining, providing a rapid, label-free alternative to conventional sectioning-based workflows. Together, these results establish TF-QPM as a scanless, high-speed platform for rapid, label-free volumetric imaging across both basic research and translational applications.

## Main

Quantitative phase microscopy (QPM) has emerged as a powerful imaging modality for characterizing cellular and tissue architecture^1,2^. By mapping cell and tissue 3D morphology through measuring optical phase retardations induced by transparent, label-free specimens, QPM offers distinct advantages over of fluorescence techniques such as confocal and multiphoton microscopy, including reduced photobleaching and phototoxicity. The resulting phase maps provide absolute optical path length information, enabling reproducible and quantitative metrics of morphology and spatiotemporal dynamics without the need of exogenous contrast agents. In transmission configurations, QPM has been widely employed to extract diverse biophysical parameters of biological specimens, including cellular dry mass^3–5^, the mechanical response of red blood cells under pathological conditions^6–8^, and tissue-level scattering^9^ and polarization signatures^10^. However, most transmission QPM intrinsically lacks of depth selectivity, unless multiple acquisitions are performed for tomographic reconstruction^11^, and it is therefore restricted to thin or relatively transparent sample, motivating the development of reflection-mode approaches for volumetric imaging in thick and scattering tissues^12^.

Reflection-mode QPM uniquely enables selectively depth-resolved imaging of thick, opaque, or highly scattering specimens in an epi-geometry, making it particularly attractive non-invasive and *in situ* tissue imaging^13–15^. Reflection-mode QPM is also intrinsically more sensitive than transmission-mode QPM to the optical path length changes^16,17^. Despite these advantages, the development of high-resolution and high-throughput 3D reflection-mode QPM remains at an earlier stage of technical development. Current strategies typically depend on sequential acquisitions or multiplexed detection, which fundamentally constrain the attainable acquisition frame rate, even though high-speed volumetric imaging is essential for resolving dynamic biological processes in real time and for delivering robust, motion-tolerant assessments in translational and clinical settings.

Point- or line-scanning implementations, such as confocal QPM^18,19^, provide optical sectioning via the confocal gating but inherently operate at reduced acquisition speeds due to their sequential acquisition nature. Full-field reflection-mode architectures offer potential for higher imaging throughput. Among these, temporal coherence-gated full-field reflection-mode QPM, often referred to as an optical coherence microscopy (OCM), has been demonstrated as a means to achieve depth-resolved imaging without lateral scanning^20–22^. In OCM, axial resolution is fundamentally determined by the temporal coherence length of the illumination, directly coupling sectioning strength to source bandwidth. For example, achieving sub-micron axial resolution at a central wavelength of 800 nm requires spectral bandwidth exceeding 280 nm, a regime typically accessible only with supercontinuum or thermal light sources. In practice, such sources often provide limited usable optical power, while supercontinuum illumination in particular introduces excess intensity noise and spectral complexity that can give rise to coherence artifacts and speckles, degrading phase stability and constraining achievable imaging throughput^12,23^.

Alternative strategies based on wavelength or phase-modulation^14,19^, rotating diffusers^24,25^, galvo scanners^26^, or digital micromirror devices^27^ have also been explored for full-field reflection-mode QPM. However, these approaches rely on additional modulation or scanning elements that increases system complexity and impose an upper bound on imaging speed. As a result, reflection-mode 3D QPM with sub-micron axial resolution has yet to exceed 200 fps, underscoring the need for new strategies to achieve truly high-speed volumetric imaging. The proposed temporal-focusing QPM is fundamentally different from OCM: whereas OCM achieves axial resolution through temporal-coherence gating, our temporal-focusing approach exploits depth-dependent pulse broadening to induce temporal decorrelation, thereby fully utilizing the numerical aperture (NA) of the objective and enabling high axial resolution without requiring ultrabroadband light sources.

Temporal focusing-based imaging was first introduced for scanless, wide-field, depth-resolved multiphoton fluorescence microscopy^28–31^ by confining light in time rather than solely relying on tight spatial focusing^32,33^. In this approach, a dispersive optical element placed in a plane conjugated to the objective focal plane introduces depth-dependent temporal pulse broadening, resulting in axially confined multiphoton excitation. While temporal focusing has been widely adopted in incoherent nonlinear fluorescence microscopy for rapid 3D imaging, its application to coherent, phase-sensitive imaging has remained largely unexplored. Here, we demonstrate that temporal focusing can be harnessed to enable single-shot, reflection-mode 3D QPM with sub-micron optical sectioning and kHz acquisition frame rates.

The TF-QPM framework eliminates the need for mechanical scanning or multiplexed measurements and requires neither supercontinuum nor thermal light sources, resulting in a compact and robust optical architecture. Using only modest spectral bandwidth, we achieve lateral and axial resolution of 402 nm and 920 nm, respectively, with reduced susceptibility to speckle noise owing to multi-angle and spectrally independent illuminations^31^. We demonstrate that phase-sensitive TF-QPM unlocks 3D particle-tracking microrheology with sub-nanometer axial displacement sensitivity—surpassing the limits of existing technologies—and extends seamlessly to rapid particle image velocimetry with dramatically reduced axial uncertainty at rates exceeding 3,780 Hz, over an order of magnitude faster than existing full-field 3D QPM approaches and establishing a new paradigm for high-fidelity, volumetric measurement in dynamic microscale systems. Finally, we show that TF-QPM supports high-throughput volumetric tissue imaging with resolution comparable to whole-slide histopathology, enabling pixel-to-pixel virtual staining in intact samples. Together, these results establish fast 3D QPM as a powerful, versatile, and label-free imaging platform for rapid volumetric imaging across both fundamental research and translational applications.

## Results

### Principle of temporal-focusing QPM

The TF-QPM imaging platform is based on a Linnik interferometer configuration, employing two identical high-numerical-aperture (NA) water-immersion objectives (Olympus LUMPLFLN 60XW, NA = 1.0) in both the reference and sample arms (Fig. 1a). The spectral components of a pulsed laser source (NKT Photonics SuperK; 5 ps pulse width, 800 nm nominal central wavelength, 40 nm spectral bandwidth after bandpass filtering) are spatially dispersed by a high-density grating and temporally recombined only at the physical focal plane of the objectives. Although the experimental demonstrations reported here use a spectrally filtered supercontinuum source, the TF-QPM is in principle compatible with any pulsed light source providing sufficient spectral bandwidth to achieve desired field-of-view (FOV, see Discussion Section IV for details).

**Fig. 1.**
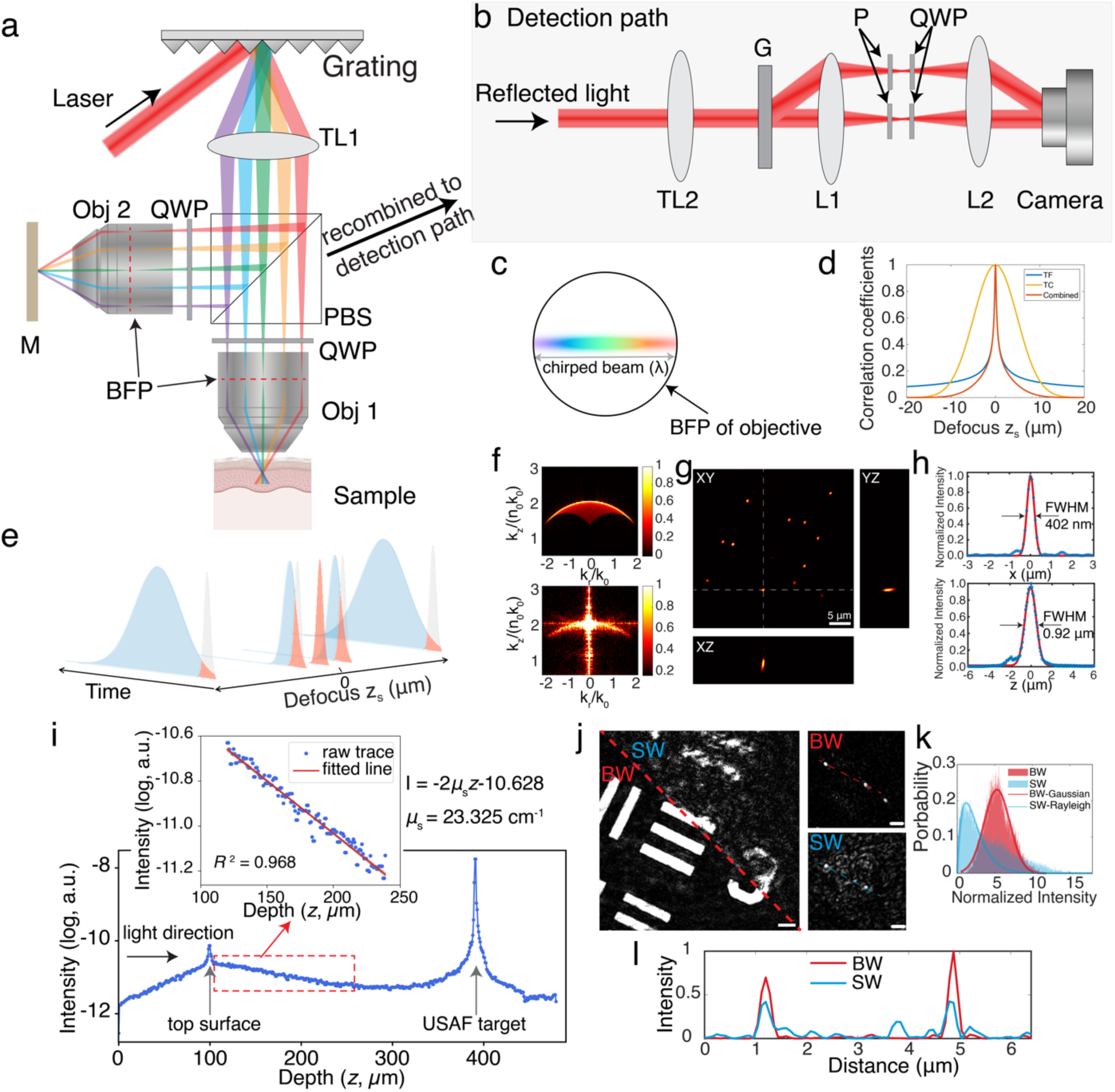
Principle and characterization of TF-QPM. **a**, setup of illumination path in imaging system. TL: tube lens, Obj: objective, BFP: back focal plane, QWP: quarter-wave plate, PBS: polarization beam splitter. **b**, detection path of recombined backscattered signals from sample and reflected signal from reference path. Two beams are recombined and collimated by TL2. The zero- and first-order of diffracted beam after grating G is then focused where a polarizer P and QWP are used to separate the beams from two paths. The signals are then interference with each other at camera plane generating interferogram. **c**, chirped beam distribution at the BFP of objectives. **d**, simulated cross-registration of interfered signals between sample and reference path. The simulation were conducted on different mechanisms, including temporal focusing (TF), temporal coherence gating (TC) and combined two mechanisms. **e**, schematic plot shows the cross-correlation between pulses from sample and reference paths. Shaded gray and blue areas indicate the pulses from sample and reference paths, respectively. Shaded red areas indicate the overlap (cross-correlated) area between two paths. **f**, simulated (top) and measured (bottom) *k*_*z*_*k*_*r*_ - projected magnitude map of optical transfer function of TF-QPM in spatial-frequency domain. **g**, measured PSF of 300-nm diameter gold nanoparticles and **h**, measured lateral and axial resolution with fitting curve. **i**, reconstructed depth-dependent backscattering intensity and fitted scattering coefficient of scattering phantom. **j**, reconstructed intensity images of USAF target (left, scale bar 5 *μm*) and 500-nm diameter gold nanoparticles (right, scale bar 2 *μm*), both are beneath a scattering phantom, by using TF-QPM (denoted as BW in the figure) and conventional full-field phase imaging (denoted as SW in the figure). **k**, histograms with fitted statistic distributions and **l**, line profile indicated by dashed curves, of measured intensity images at i.

To fully utilize the NA of the objectives, the tube lens (TL1) was selected such that the chirped beam overfills the back pupil planes of both objectives along the dispersion direction (Fig. 1c). This configuration produces a depth-dependent temporal broadening of the pulses, with the shortest pulse width confined to the conjugate focal plane of the grating. Depth-encoded backscattered signals from the sample arm are then recombined with the reference-arm reflections from a mirror positioned at the focal plane. Off-axis holography is used to record a single-shot interferogram, from which the complex optical field (amplitude and phase) information is recovered^34^ (Fig. 1b).

In TF-QPM, the temporal focusing introduces depth-dependent pulse broadening in the backscattered field from the sample, while the reference field remains temporally compressed at the focal plane. As a results, the detected intensity *I*(*r*, *z*_*s*_) of interference field at the camera plane is governed by the temporal cross-correlation between the depth-encoded backscattered fields from the sample arm *U*_*bs*_(*r*, *z*_s_)*e*^−*α*(*zs*)*t*2^ and reflection field from a mirror positioned at the objective focal plane *U*_*R*_*e*^−*α*(*zs*|*zs*=0)*t*2^ (see Supplementary Information Section I for detail discussion). Specifically, the interference intensity can be expressed as:

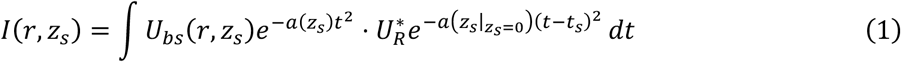

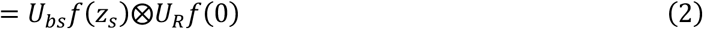

where *f*(*z*_s_) = *e*^−*α*(*zs*)*t*2^ describes the Gaussian temporal envelope of the pulse at axial position *z*_*s*_, and ⊗ denotes convolution. *r* denotes the lateral dimension coordinates. *U*_*R*_and *U*_*bs*_ represents the complex optical fields from the reference and sample arms, respectively. *z*_*s*_ denotes the defocusing distance with respect to the focal plane in the sample arm, and *t*_*s*_ = *n*_*m*_*z*_*s*_/*c* is the relative time delay between the two signals from the sample and reference arms. The parameter *α*(*z*_*s*_) = 2*ln*2/*τ*^2^(*z*_*s*_) represents the axial-dependence of pulse broadening due to temporal focusing and *τ*(*z*_*s*_) is the pulse width at depth *z*_*s*_. The pulse width *τ*(*z*_*s*_) is minimized at the focal plane where the temporal overlap (cross-correlation) between the sample and reference fields is maximized. With backscattered light from defocused depths, the increasing temporal decorrelation between the two fields reduces the interference signals, thereby providing intrinsic depth-resolved image contrast (Fig. 1e).

Considering the experimental configuration in which an axially scanning mirror is placed in the sample arm while the reference mirror remains fixed at the focal plane of the reference arm, the interference signal detected at camera, i.e., normalized cross-correlation between the two fields, can be written as (see Supplementary Information Section II):

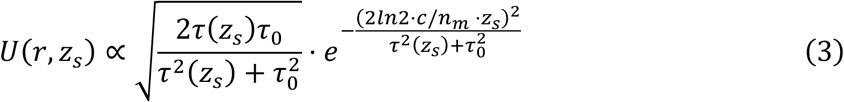

where *c* is the light speed and *n*_*m*_is the refractive index of medium. This expression shows that axial response is governed by the temporal correlation between the sample and reference pulses and therefore depends on the depth-dependent pulse width *τ*_*s*_(*z*). In TF-QPM, the axial pulse broadening is determined by the NA of the objective^31^ (see Supplementary Section II) can be approximately as:

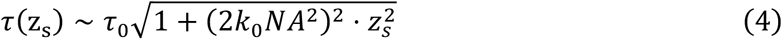

where 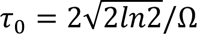 is the pulse width at focal plane, Ω is the full-width half-maximum (FWHM) of the pulse spectrum of the pulse and *k*_0_ is the wavevector of central wavevector. Stronger temporal focusing is therefore achieved with increasing NA of objective. Notably, this axial sectioning mechanism is largely independent of the source width, in contrast to conventional low-coherence microscopy^35^ (see Discussion). From the calculated pulse cross-correlation, an axial resolution of approximately 1.0 *μm* is predicted for an objective with a NA of 1.0 (Fig. 1d).

### Optical transfer function of TF-QPM

To obtain the optical transfer function (OTF) of TF-QPM, we rigorously model the pulsed light propagation in the specimen by solving the wave equation under a modified first-order born approximation in spatial-frequency domain. When both sample and reference objectives are illuminated with a chirped pulse laser beam, the 3D OTF Γ(*k*_*x*_, *k*_*y*_, *z*_*s*_) can be expressed as a function of lateral spatial frequency ***k***_***r***_ = (*k*_*x*_, *k*_*y*_), axial defocus *z*_*s*_, pupil function *P*(*k*_*x*_, *k*_*y*_, *ω*) and temporal coherence function *S*(*ω*) = *e*^−(*ω*–*ω*0)2/(Δ*ω*)2^, integrated over the entire source spectrum Δ*ω* (see Supplementary Information Section I for details):

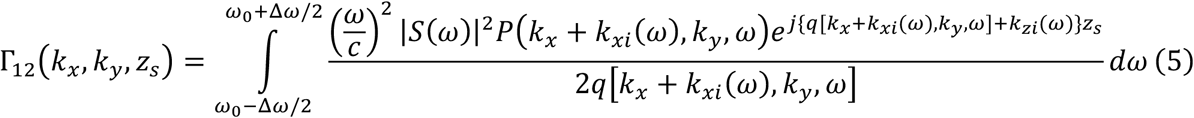

Here,

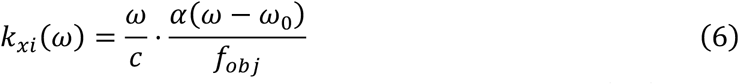

describes the spatially chirped at back focal plane (BFP) of the objectives. In our case, (*k*_*xi*_)_*mαx*_ = *k*_0_*NA* where *k*_0_ is the wavevector of central spectrum. *f*_*ob*j_ is the focal length of objectives and *α* is the dispersion parameter, determined by the grating period and focal length of the tube lens. In our implementation, the chirped beam overfills the objective BFP (diameter *d*), yielding:

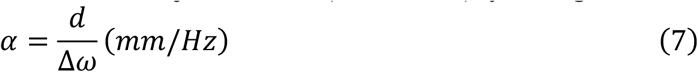

which is evaluated to be *α* = 5.1 × 10^−14^ *mm*/*Hz*. Compared with conventional temporal focusing multiphoton microscopy with transform-limited femtosecond laser pulse^29^, the broader spectral width employed here leads to a smaller effective dispersion parameter. The 3D OTF can be numerically simulated based on the Eq. (5). In the special case ***k***_***r***_ = 0, where the sample is a simple infinitely thin plate translating along light propagation direction, the axial resolution of our system can be obtained as:

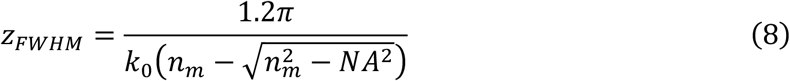

This corresponds to an axial resolution of *z*_*FWH*M_ ∼ 1.06 *μm* for NA = 1.0, with the immersion medium refraction index *n*_*m*_ = 1.33 and central wavelength *λ*_0_ = 800 *nm*. Notably, both rigorous OTF analysis and pulse-correlation theory indicate that the axial resolution of TF-QPM scales as 1/(*k*_0_*NA*^2^), consistent with most reflection-mode 3D imaging systems^12^.

### Experimental validation of OTF and self-healing behavior

To characterize system performance, we imaged 300-nm gold nanoparticles sparsely embedded in agarose gel. The simulated OTF (Fig. 1f, top) closely matches the measured OTF (Fig. 1f, bottom). In temporal focusing, the chirped pulses fully illuminate only a narrow line of the BFP, resulting in reduced OTF magnitude near the center of the frequency cone compared to its periphery, as observed in both simulation and experiment. The measured lateral and axial resolution are 402 nm and 0.92 *μm*, respectively, in good agreement with simulation results (Fig. 1g and Fig. 1h).

An additional advantage of TF-QPM is its inherent “self-healing” property^31^, which enhances beam robustness to scattering and preserves spatial resolution compared to conventional full-field illumination (a single-spot is focused at the BFP of objective and generate a single-angle full-field illumination at sample space). To demonstrate this, we imaged a standard reflective USAF target position beneath a scattering phantom with a measured scattering coefficient of *μ*_*s*_ = 23.3 *cm*^−1^ (scattering length *l*_*s*_ = 429 *μm*). At an imaging depth into the phantom of approximately 225 *μm* (∼ 0.5*l*_*s*_), TF-QPM exhibited improved signal-to-noise ratio (SNR) and reduced resolution degradation compared with conventional single-angle full-field QPM (Fig. 1i, l). Furthermore, the intensity histogram of reconstructed TF-QPM intensity images followed a Gaussian distribution, whereas single-wavelength full-field QPM exhibited a Rayleigh distribution (Fig. 1k), indicating reduced speckle noise in TF-QPM^36^. The self-healing behavior is attributed to the fact that different spectral components are incident at different angles, generating uncorrelated speckle patterns during propagation to the focal plane, which are subsequently averaged at the focal plane to yield a speckle-reduced image^31^ (see more discussions in Supplementary Information Section III).

### Phase-sensitive imaging with nanometer height sensitivity at kHz frame rate

To validate the phase-sensitive optical sectioning capability of TF-QPM, we first imaged a commercial phase calibration target (Benchmark Technologies; Fig. 2a). The periodic structures were clearly resolved in both reconstructed intensity and phase images, with the phase data enabling accurate height quantification beyond axial resolution limit. The extracted height profile of the target (Fig. 2b) yielded a mean height value of 102.51 ± 2.36 nm, in close agreement with the 104.2 nm measured by using atomic force microscopy.

**Fig. 2.**
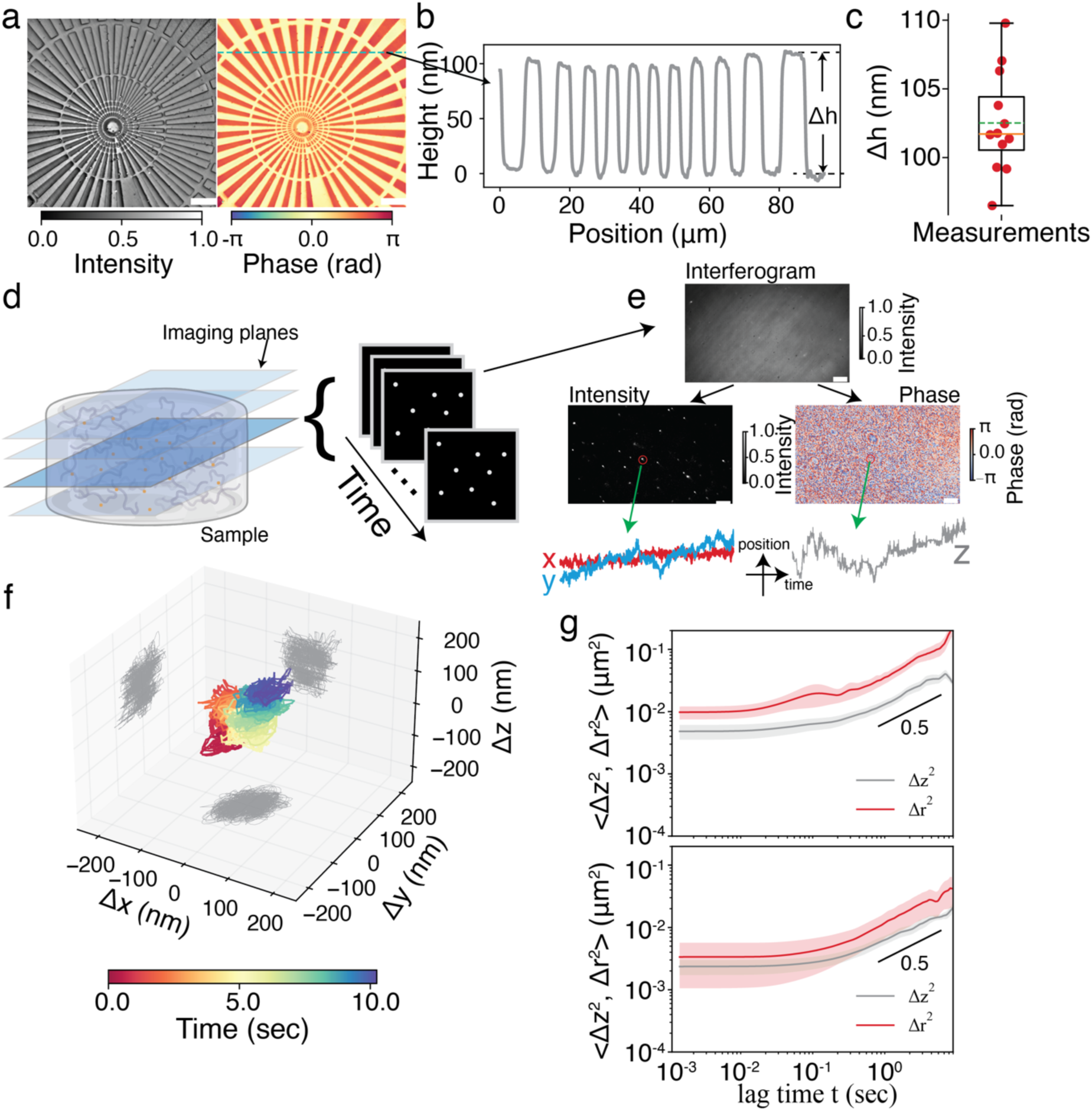
Phase-sensitive imaging enabled by TF-QPM. **a**, reconstructed intensity (left) and phase (right) images of standard star phase target (scale bar 10 *μm*). **b**, height profile (green dashed line in a) and **c**, statistic distribution of measured by phase imaging from right of a. the measured phase was converted to height by Δℎ = *Φ* ⋅ *λ*/4*πn*_*m*_where *Φ* is the phase, *λ* = 800 nm is the center wavelength of light source and *n*_*m*_= 1.33 is the refractive index of medium. **d**, schematic illustration of imaging beads fluctuation suspended in agarose gel at multiple planes. The time-lapse imaging was conducted sequentially at each plane. The obtained interferograms were reconstructed to obtain intensity images for lateral (xy) tracking and phase images for axial tracking (scale bar 5 *μm*). **f**, representative 3D trajectory of bead fluctuation. **g**, mean square displacement (〈Δ*r*^2^〉 for lateral tracking, 〈Δ*z*^2^〉 for axial tracking) of nanoparticles versus time lag measured in different concentration agarose gel (0.07% w/v top, and 1.2% w/v bottom).

High-throughput imaging with nanometer-scale axial sensitivity enables applications that are difficult to access with existing 3D imaging modalities, particularly in biological and soft-matter systems. Here, we demonstrate two representative capabilities enabled by TF-QPM: 3D particle-tracking microrheology and high-speed particle image velocimetry (PIV), both with nanometric positional sensitivity. While particle tracking and PIV are well established technologies, high-speed volumetric tracking with near isotropic nanometer-scale sensitivity remains highly challenging, and our approach provides unprecedented capability for probing the biomechanics of anisotropic materials and bio-systems^37–40^ (see comparison between our approach and existing PIV techniques in Supplementary Information Section V).

3D particle tracking microrheology provides non-invasive, spatially resolved measurements of microscale viscoelastic properties in complex and heterogeneous biological environments, overcoming the limitations of bulk rheology and enabling mechanical characterization of 3D cell– matrix systems that are critical for understanding disease progression and tissue function^41,42^. However, most existing implementation rely on 2D wide-field imaging with limited axial sensitivity (in most optical system, axial resolution is usually a few times worse than lateral resolution), implicitly assuming mechanical isotropy – an assumption that often fails in biological cells and tissue environments. TF-QPM demonstrates high axial displacement sensitivity (∼2 nm, equivalent to 40 mrad phase sensitivity), enabling full 3D particle-tracking microrheology. We tracked 500 nm diameter polystyrene particles embedded in agarose gels (0.07% w/v and 1.2% w/v) over 10 sec sequences at four axial planes with frame rate of 780 Hz (Fig. 2d). Lateral particle motion was extracted from the reconstructed intensity while the axial displacements were obtained from the phase measurements (Fig. 2e). In softer gel, particle trajectories showed random 3D motion with ∼100 nm amplitude (Fig. 2f). The mean square displacements (〈Δ*r*^2^〉, 〈Δ*z*^2^〉) scaling followed power-law in both gels but with smaller scaling exponents for the stiffer gel (0.51 vs 0.57, Fig. 2g). Notably, in stiffer gel, the lateral mean square displacement 〈Δ*r*^2^〉 showed greater variability than axial mean square displacement 〈Δ*z*^2^〉, highlighting an enhanced robustness of phase-based axial tracking over intensity-based lateral tracking^43,44^. Although these proof-of-concept demonstrations focus on isotropic systems, the combination of axial displacement sensitivity and high imaging speed of this approach can be readily applied to the study of complex, heterogeneous 3D biological systems^37,45^.

We further demonstrate the capability of TF-QPM for high-speed 3D particle imaging velocimetry (PIV). PIV is a non-intrusive optical technique that provides instantaneous, spatially resolved velocity fields and has become a widely used tool for quantitative flow characterization across fundamental fluid mechanics and a wide range of industrial and biomedical applications^46^. In volumetric PIV, high axial resolution critical to minimize depth-induced correlation errors and to accurately reconstruct 3D flow fields^47^. Existing reflection-mode PIV implementation face a trade-off between axial resolution and imaging speed. TF-QPM overcomes this limitation by providing high-precision 3D particle localization (nanometer-scale axial localization derived from the phase measurements with sub-diffraction lateral localization achieved through intensity-based centroiding) at kHz frame rate without multiplexing or mechanical scanning. TF-QPM therefore can open up the potential of *in vivo* quantitative PIV in biological systems (e.g., blood flow^48^) for diseases diagnostics and other complex system that is inaccessible by the transmission-mode PIV^38^. We tracked nanoparticles in flow at frame rate of 3.7 kHz with temporal projection shown in Fig. 3a. The high axial resolution of TF-QPM suppresses contributions from out-of-focus particles, reducing velocity uncertainty. The measured trajectories of nanoparticles revealed consistent flow profiles, with distinguishable mean velocities of 121.0 ± 6.0 *μm*/*s* in water versus 102.0 ± 6.1 *μm*/*s* in 10% w/w glycerol.

**Fig. 3.**
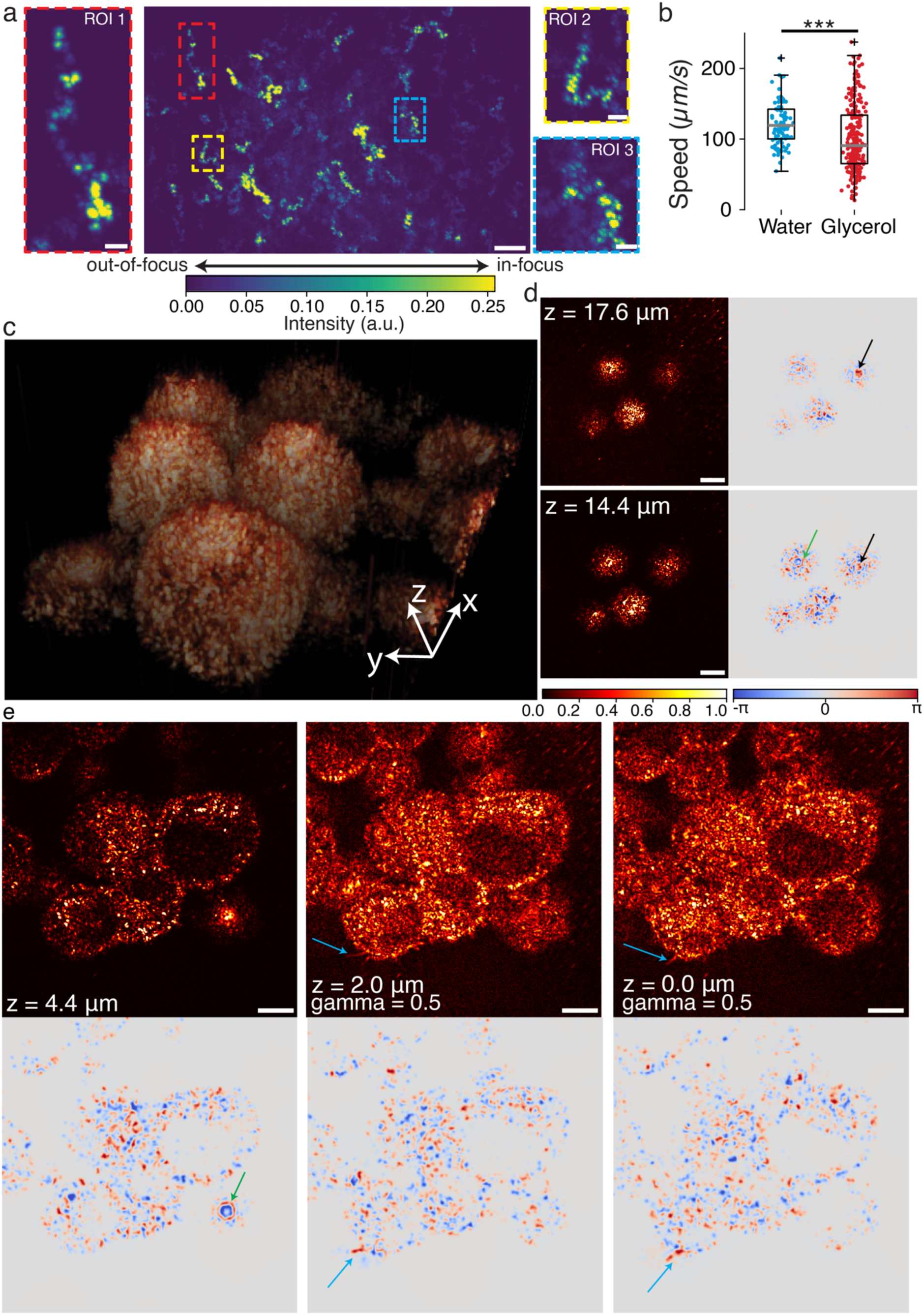
Rapid optical sectioning imaging enabled by TF-QPM. **a**, time-lapse projected intensity image of 300-nm diameter nanoparticles suspended in water (scale bar 5 *μm*). The time-lapse intensity images were reconstructed from acquired interferograms at a single plane over 10 seconds. The coverslip of petri-dish was cut to open a small slit to trigger the flowing of nanoparticles in solutions. **b**, measured speed distributions of nanoparticles in both water and 10% w/w glycerol solutions by using the same petri-dish. **c**, volumetric rendering view of multi-cellular systems (MDA-MB-435S) with field of view 90 *μm* × 90 *μm* × 23 *μm* along xyz dimensions. **d-e**, reconstructed intensity (left at d and top at e) and phase (right at d and bottom at e) images of cells at different depths. The intensity images at middle and right columns of e were gamma-corrected by a factor of 0.5 to improve image contrast. Scale bars are 10 *μm*.

### High-throughput virtual staining of biological cells and tissues at subcellular resolution

We next applied TF-QPM to 3D morphological imaging of melanoma cells (MDA-MB-435S, Fig. 3c). The reconstructed 3D volumes captured whole-cell architecture (Fig. 3c) and revealed subcellular features across multiple depths, including nuclei and nuclear boundaries (Fig. 3d black arrows). Depth-resolved phase imaging distinguished plasms and nuclear membranes, separated by ∼ 3 *μm*, with markedly enhanced contrast compared with intensity images (Fig. 3d right column). Multiple filopodia were observed near the surface of cells (light-blue arrows in Fig. 3e). Notably, the nuclear contours were clearly visualized in the phase images (green arrow in Fig. 3d-e), but were poorly discernible in corresponding intensity images, highlighting the advantage of phase-sensitive optical sectioning.

TF-QPM further enabled rapid, high-throughput 3D imaging of Masson Trichrome–stained human colon adenocarcinoma biopsy specimens (iHisto Inc., RA99-34305) over millimeter-scale FOVs at subcellular resolution within 2 minutes. Large-area images (1.4 mm × 1.6 mm) were acquired by stitching multiple 3D stacks (70 μm × 80 μm × 100 μm each) collected at 10 volumes/s (0.5 μm axial steps) using a piezoelectric scanner and an XY motorized stage. The representative acquired single volume of tissue block is shown in Fig. 4b, illustrating different tissue features at different depths. The maximum intensity projection (MIP) of the reconstructed volumes (Fig. 4c) closely matched whole-slide histopathology and revealed hallmark features of colon adenocarcinoma, including irregular infiltrative glandular architectures, luminal structures (orange arrows) and goblet cell features (green arrows) (Fig. 4d-g). Additional diagnostic features, including distinct nuclei morphology (yellow arrows), collagen-rich fibrosis and muscle fibers (purple arrows), were clearly visualized, along with regions of dirty necrosis (Fig. 4f, light-blue arrows in Fig. 4f), a characteristic indicator of ischemia associated with rapid tumor growth in primary colorectal adenocarcinoma^49,50^.

**Fig. 4.**
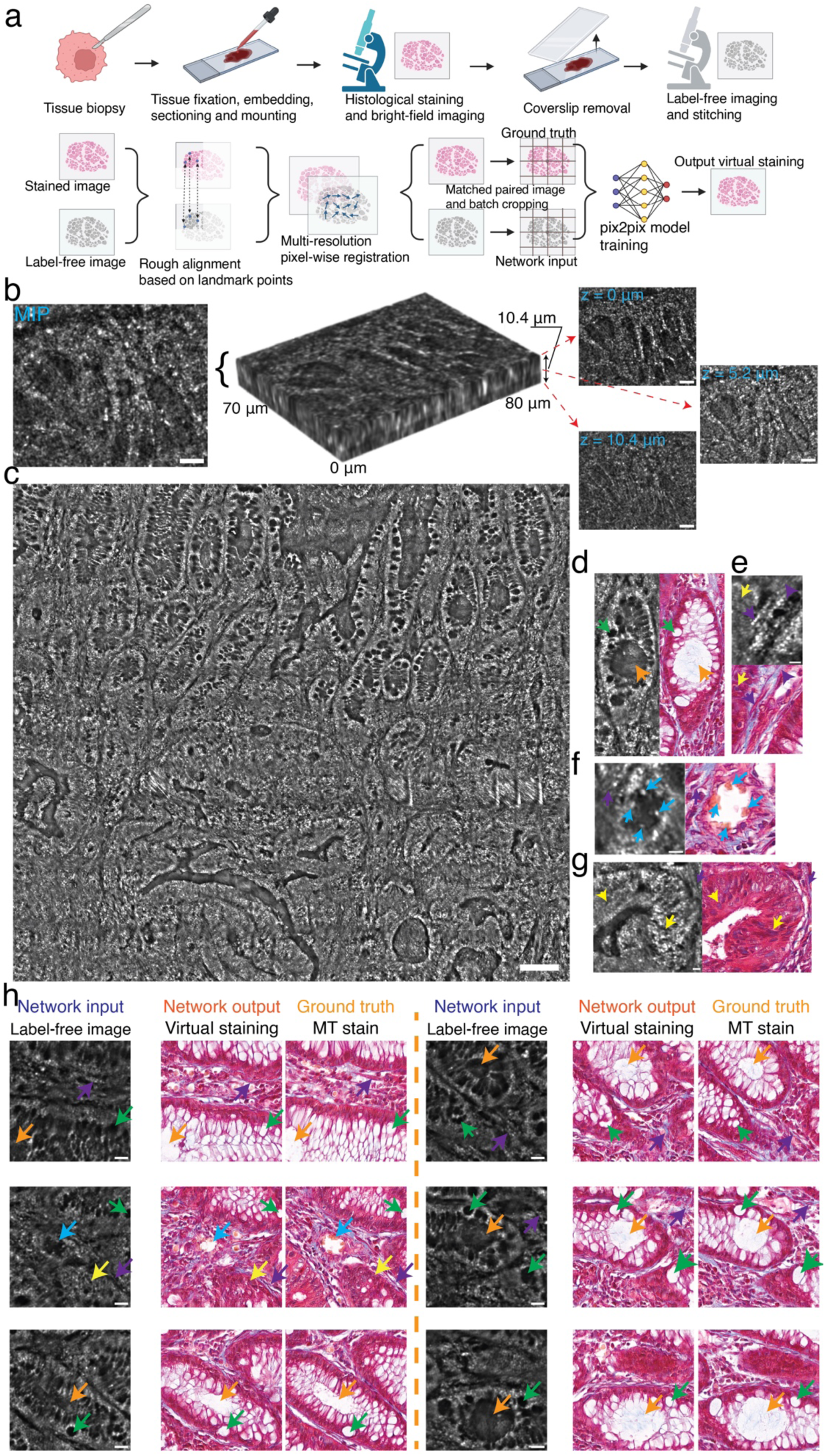
High-throughput tissue imaging at sub-cellular resolution enabled by TF-QPM. **a**, sample preparation (top row) and cross-registration (bottom row) for immunostaining and label-free images of human colon adenocarcinoma tissue. **b**, representative volumetric intensity images (middle), MIP (left) and images at different depths (right, scale bar 40 *μm*) of colon cancer tissues measured by TF-QPM. **c**, stitched MIP images across FOV of 1.4 mm x 1.6 mm with pixel number of 15217 x 17391 from reconstructed intensity images by TF-QPM (scale bar 100 *μm*). **d-g**, selective representative features of label-free images (left columns) from c (scale bar 40 *μm*), compared to paired immunostaining images (right columns). **h**, virtual staining results and comparison to ground-truth immunostaining images (scale bar 40 *μm*).

We further demonstrate that the high spatial resolution of TF-QPM enables pixel-accurate co-registration between label-free images and their stained histopathology counterparts, facilitating virtual staining at same scale (Fig. 4a, bottom row). A pixel-to-pixel, deep-learning colorization network^51,52^ was trained using paired TF-QPM images and Masson trichrome-stained ground truth (Methods). Representative label-free inputs, virtual staining, and ground truth are shown in Fig. 4h. Hallmark features of colon adenocarcinoma are observed, including irregular granular architectures (orange and green arrows), infiltrating nuclei (yellow arrows), and dirty necrosis (light-blue arrows). The average structural similarity index reaches ∼ 0.76 ± 0.029, comparable to the performance reported in state-of-the-art virtual staining approaches^53,54^.

Discrepancies between virtual stains and ground truth primarily arise from the strict pixel-wise correspondence required for supervised training. In our workflow, fixed specimens were immersed overnight in a dissolvable medium to remove the coverslip for suppressing the interface reflections, introducing local tissue distortion that degrade sub-cellular co-registration. Registration-robust learning strategies with unpaired or weakly supervised frameworks, as well as incorporation of confocal gating in the imaging system could further improve the virtual staining performance^18,27^. Moreover, the ground-truth histopathology images were acquired in transmission mode, which predominantly preserves low-frequency axial content. Since TF-QPM captures high-frequency axial features in reflection mode, training with reflection-mode ground truth, such as light-sheet and confocal images, may further enhance virtual staining accuracy. Nevertheless, these proof-of-concept results demonstrate the potential of TF-QPM as a rapid, label-free platform for high-throughput virtual staining in translational and *in situ* pathology applications.

Reflection-mode label-free virtual staining offers a transformative path toward rapid, nondestructive, slide-free histology by directly obtaining diagnostic tissue contrast from intact, opaque specimens, creating a foundation for true *in vivo* histopathology without fixation, sectioning, or chemical staining^55^. Beyond transmission-mode approaches (e.g., autofluorescence), reflection-mode approach is complementary and provides additional depth selectivity through thick opaque tissues and unlocks translational in vivo applicability, yet current implementations remain limited by insufficient 3D resolution and acquisition speed that restrict practical deployment^52,55–57^. Our proposed TF-QPM overcomes these barriers by combining high-speed sampling robust to motion, sub-cellular 3D resolution capable of revealing fine morphological hallmarks, and reduced speckle noises with enhanced contrast to elevate diagnostic fidelity. Moreover, its architecture can be seamlessly adapted into a swept-source endoscopic platforms^58^, enabling real-time, high-resolution intraluminal imaging of gastrointestinal and vascular tissues and advancing minimally invasive, sub-cellular-scale early disease detection^59^.

## Discussion

We have demonstrated a full-field, reflection-mode quantitative phase microscopy platform that enables depth-resolved phase-sensitive imaging at kilohertz frame rate with sub-cellular resolution. By introducing a spatially chirped pulsed illumination, TF-QPM extends temporal focusing to coherent, full-field phase-sensitive imaging, where axial sectioning arises from the decorrelation of temporally broadened pulses and is ultimately determined by the NA of the objective. In addition to enabling optical sectioning, the spatially chirped illumination suppresses speckle and better preserves SNR in scattering media more effectively than conventional full-field implementations.

We further demonstrate depth-resolved particle-tracking microrheology and high-speed particle imaging velocimetry, achieving improved 3D displacement sensitivity and resolving features at closely spaced axial planes. At spatial scale comparable to histopathology, TF-QPM also supports high-throughput 3D imaging of cells and tissues, enabling pixel-wise virtual staining through direct cross-registration between label-free phase and stained ground truth images. The reflection-mode implementation makes our approach particularly flexible and practical for translational and *in situ* biomedical applications.

Several practical considerations currently limit system performance. First, volumetric acquisition speed is constrained by the axial actuator. Implementing remote focusing with rapid dispersion control^29,31^ could increase imaging throughput to hundreds of volumes per second while preserving phase fidelity. Second, out-of-focus background intensity can consume camera dynamic range and degrade phase sensitivity. Incorporating a line-scanning confocal gate with a resonant scanner could reject most out-of-focus light without compromising camera speed, thereby increasing the effective dynamic range available for interference and further improving phase sensitivity^60^. Third, the current implementation of temporal focusing results in a line illumination at back pupil planes of objectives but does not fill it, which degrades the performance of optical sectioning analogous to the fluorescence case^61^. Employing line-scanning geometry or two-dimensional spectral dispersion and temporal shearing with orthogonal grating pairs could extend the pulse stretching across the entire pupil planes, improving the robustness and sectioning performance^62^.

In temporal-focusing microscopy, the maximum usable FOV is fundamentally constrained by the illumination bandwidth. Although broader spectra increases angular dispersion and enhance axial confinement, they also accelerates wavelength walk-off across the focal plane: the additional group delay variation across the field scales approximately with the square of the FOV, limiting the lateral region over which the pulse remains temporally compressed^63^. Additional geometric constrains in the relay and scan optics (see Supplementary Information Section IV) impose additional bounds on the achievable FOV. For field-of-view on the order of hundreds of micrometers, spectral bandwidths of ∼10 nm are generally sufficient, making many femtosecond light sources well suited for this design. Furthermore, because the scattered signal in TP-QPM is orders of magnitude stronger than nonlinear fluorescence processes, low-power and cost-effective femtosecond lasers are adequate for practical implementations.

Taken together, TF-QPM provides a compact, scan-free approach to rapid three-dimensional, subcellular-resolution, phase-sensitive imaging with enhanced robustness in scattering tissues. We anticipate that this capability will enable broad applications in many frontiers of biomedical imaging, including quantitative studies of 3D biophysical properties in complex biological materials and high-throughput label-free pathological analysis of clinical specimens.

## Methods

### Imaging setup

A pulsed laser (NKT Photonics SuperK, pulse width 5 ps, nominal central wavelength of 800 nm with bandwidth of 40 nm) linearly polarized at 45 degree is used for illumination. The output beam is spectrally dispersed by using a diffraction grating. In the present implementation, a digital micro-mirror device (DMD, Vialux DLP V-7001 VIS, 1024×768 pixels with pitch size of 13.7 um and maximum switching rate of 22.7 kHz) was employed as a programmable diffraction grating to facilitate optical alignment; however, a conventional one-dimensional high-density diffraction grating can be used equivalently. After spectral dispersion, different wavelength components are then focused using a tube lens (TL1) and a polarization beam splitter (PBS) to different lateral positions at the back pupil planes of both sample and reference objectives. To fully utilize the NA of the objectives along the dispersion direction, the TL1 and a subsequent telescope module (not shown in Fig. 1a) is employed such that the spectrally dispersed beam overfills the back pupil planes of the objectives along the chirped dimension. In the reference arm, a high-reflectivity mirror positioned at the focal plane of the objective provides the reference reflection.

Quarter-wave plate (QWP) oriented at 45 degrees were placed in both interferometer arms between the microscope objectives and the PBS, such that each beam undergoes a 90-degree polarization rotation upon double pass. As a result, the cross-polarized returning fields from sample and reference arms recombine at the same PBS into a common detection path (Fig. 1b), where they are collimated by a tube lens (TL2) and then diffracted by a transmission grating which is conjugate to the camera detection plane. A 4f relay in combination with crossed polarizers (*P*_∥_ and *P*_⊥_) is used to separate the signals from sample and reference arms. Two additional QWPs at 45 degrees are used to align the polarization states of the sample and reference fields, enabling their interference with a well-defined angular carrier set by the density of grating G. The raw interference pattern obtained from camera is 2D Fourier transformed to isolate the first-order diffraction term, followed by an inverse-transform operation to recover the complex optical field, from which both amplitude and phase images are obtained. The intensity images are calculated as the square of the reconstructed amplitude images.

### Histology and imaging

Human colon adenocarcinoma tissue was obtained by biopsy and immediately fixed in 4% paraformaldehyde (PFA) in PBS for 16 h at 4 °C. Following fixation, tissue was cryoprotected in 30% (w/v) sucrose in PBS at 4 °C until fully saturated, embedded in optimal cutting temperature (OCT) compound, and stored at −80 °C until sectioning. Fixed-frozen samples were cryosectioned at the desired thickness of 10 *μm* using a cryostat, mounted on glass slides, and stained with Masson’s Trichrome staining to differentiate tissue components, with collagen visualized in blue, cytoplasm in red, and nuclei in dark purple. Stained tissue slides were imaged using a whole-slide scanner equipped with a 20x objective lens (NA = 0.75), producing high-resolution digital images for histopathological analysis. All the tissue staining and immunofluorescence imaging were performed commercially by iHisto (https://www.ihisto.io/).

### Phase imaging of histology specimen

Fixed and stained human adenocarcinoma colon biopsy specimens (iHisto Inc. RA99-34305) were immersed into xylene solution overnight to remove the coverslip prior to phase imaging. For large FOV, images were acquired in a 2D grid pattern by sequentially imaging multiple adjacent regions using a XY motorized stage. At each lateral position, a full 3D image stack with volume of 70 *μm* × 80 *μm* × 100 *μm* was recorded. Imaging acquisition was performed at a camera frame rate of 1 kHz, chosen to balance available laser power and SNR. For axial scanning, the tissue sections (<0.2 g) were mounted on a piezoelectric objective scanner (PI P-725.4CD) for axial scanning, achieving 10 volumes per second with an axial stepping size of 0.5 *μm*.

### Cell culture

MDA-MB-435S cells were cultured under aseptic conditions in Dulbecco’s Modified Eagle Medium (DMEM; ATCC, 30-2002) containing 4 mM L-glutamine, 4500 mg/L glucose, 1 mM sodium pyruvate, and 1500 mg/L sodium bicarbonate, supplemented with 10% fetal bovine serum (FBS; ATCC, 30-2021). Cells were maintained in T75 flasks at 37 °C in a humidified incubator with 5% CO₂. The culture medium was replaced every 1-3 days. Cells were subcultured every 3-7 days by trypsinization to prevent overconfluence; excess cells were cryopreserved in 95% complete medium supplemented with 5% dimethyl sulfoxide (DMSO). For imaging preparation, cells were trypsinized, centrifuged at 150 x g for 10 minutes, and seeded into a petri dish. Cells were cultured for a minimum of 12 hours prior to imaging to ensure robust attachment.

### Phantoms

For point spread function (PSF) characterization, 50 µL suspension of 300-nm diameter gold nanoparticle solutions (Sigma-Aldrich, #742082) were dispersed in 2% w/w agarose gel in water, yielding total volume of ∼ 2 mL. For scattering phantom preparations, 0.4 mL suspension of 5 wt% titanium dioxide (TiO_2_) with a nominal diameter of 900 nm (Sigma-Aldrich, #914606) was dispersed in 2% w/w agarose gel in water to a total volume of 5 mL.

### Virtual staining model

Tile-stitched maximum-intensity-projection volumes from label-free TF-QPM were downsampled and interpolated to match the pixel size of the corresponding immunostained tissue images (0.35 µm per pixel). The label-free images were initially aligned to the stained ground truth images by selecting multiple anatomically corresponding landmarks and applying a rigid transform (rotation and translation) using BigWarp^64^ in Fiji/ImageJ^65^. To further refine this coarse alignment, a multi-resolution, intensity-based pixel-wise registration was performed, yielding sub-pixel correspondence between two modalities. The resulting co-registered fields of view (4000 × 4570 pixels) were randomly cropped into 512 × 512 patches for model training and testing, with data augmentation achieved through random in-plane translations. Virtual staining was performed by using a pix2pix conditional generative adversarial network (GAN) with a unet256 generator architecture, an initial weight scaling (gain) of 0.02, and a learning rate of 2 × 10^-4^. A PatchGAN discriminator with four downsampling layers was employed. Trained models were subsequently applied to held-out images to generate virtual staining outputs.

## Data availability

The main data that support the findings of this study are available in this Article.

## Acknowledgements

All authors acknowledge support from National Institutes of Health (NIH) grant R21GM140613. Y.L., Z.Y. and P.T.C.S. further acknowledge support from NIH grant 5-P41-EB015871-27, R01HL121386, the Hamamatsu Corporation, and MIT-Fujikura Partnership Fund. P.T.C.S. further acknowledges support from NIH grant R01GM160726.

## Author information

These authors contributed equally: Xiang Zhang, Rebecca E Zubajlo

Authors and Affiliations

**Laser Biomedical Research Center, G. R. Harrison Spectroscopy Laboratory, Massachusetts Institute of Technology, Cambridge, MA, 02139, USA**

Yuechuan Lin, Zahid Yaqoob, Peter T.C. So

**Department of Mechanical Engineering, Massachusetts Institute of Technology, Cambridge, MA, 02139, USA**

Yuechuan Lin, Xiang Zhang, Rebecca E Zubajlo, Brian W. Anthony, Peter T.C. So

**Institute for Medical Engineering and Science, Massachusetts Institute of Technology, Cambridge, MA, 02139, USA**

Xiang Zhang, Rebecca E Zubajlo, Brian W. Anthony

**Neurophotonics Center, Boston University, Boston, Massachusetts, 02215, USA**

Zahid Yaqoob

**Department of Biomedical Engineering, Boston University, Boston, Massachusetts, 02215, USA**

Zahid Yaqoob

**MIT.nano, Massachusetts Institute of Technology, Cambridge, MA 02139, USA**

Brian W. Anthony

**Department of Biological Engineering, Massachusetts Institute of Technology, Cambridge, MA, 02139, USA**

Peter T.C. So

## Contributions

Y.L. and P.S. conceived the idea. Y.L. designed and carried out the experiments, conducted simulation and analyzed all the data. Y.L., X.Z. and R.Z. performed the cell imaging experiments and analyzed the imaging data. X.Z. and R.Z. prepared the cell and tissue samples for imaging. Y.L. and P.S. drafted the manuscript with inputs from all authors. P.S., Z.Y. and B.A. supervised the whole project.

## Corresponding authors

Corresponds to Peter T.C. So, Zahid Yaqoob, or Brian W. Anthony.

## Ethics declarations

### Competing interests

The authors declare no competing interests.

## Notes

### Competing Interest Statement

The authors have declared no competing interest.

