## Supplementary information for "Scanless temporal focusing enables high-speed three-dimensional quantitative phase microscopy"

#### Section I: Optical transfer function via wave equation

With dispersed beam illumination over back focal plane, it leads to:

$$U_i(x_i, y_i, \omega; z = 0) = S(\omega)\delta[x - \alpha(\omega - \omega_0)]\delta(y) \tag{1}$$

The Fourier transform of this field results in illumination fields at $\boldsymbol{k}$-space inside sample:

$$U_i\big(k_{xi}, k_{yi}, \omega; z = 0\big) = S(\omega)e^{-jk_{xi}\alpha(\omega-\omega_0)} \tag{2}$$

If considering electric field over a distance $z$, it will be represented (with angular spectrum method propagation kernel):

$$\widetilde{U_i}\big(k_{xi}, k_{yi}, \omega; z\big) = S(\omega)e^{-jk_{xi}\alpha(\omega-\omega_0)}e^{jk_{zi}z} \tag{3}$$

According to the paired coordinates between Fourier planes, the $k_x$ is then defined by the illumination wavelength component locations at back aperture of objective. It is defined as:

$$k_{xi} = 2\pi \cdot \frac{\alpha(\omega - \omega_0)}{\lambda f_0} \tag{4}$$

that's:

$$k_{xi} = \frac{\omega}{c} \cdot \frac{\alpha(\omega-\omega_0)}{f_0} \tag{5}$$

where $f_0$ is the focal length of the objective. The maximum of $\alpha(\omega - \omega_0)$ is $d/2$, which is half of the pupil aperture of the objective. To be noticed, $(k_{xi})_{max} = 2\pi \cdot \frac{d}{2\lambda f_0} = k_i NA$, with $k_i = \omega/c$. As defined, it will then determine the propagation component as:

$$k_{zi} = \sqrt{\left(\frac{n_m\omega}{c}\right)^2 - k_{xi}^2} \tag{6}$$

This Eq.(3) can be equivalently written as:

$$\widetilde{U}_i(\boldsymbol{k_{ri}}, z, \omega) = S(\omega)A(\boldsymbol{k_{ri}}, \omega)e^{jk_{zi}z} \tag{7}$$

with $A(\boldsymbol{k_{ri}}, \omega) = e^{-jk_{xi}\alpha(\omega-\omega_0)}$. This form of equation is very useful when dealing with wave equation. Or in a more general way, it can be re-written as:

$$\widetilde{U}_i(\boldsymbol{k_{ri}}, z, \omega) = S(\omega)A(\boldsymbol{k_r})\delta(\boldsymbol{k} - \boldsymbol{k_i}) \tag{8}$$

with

$$\delta(\boldsymbol{k} - \boldsymbol{k_i}) = \delta(\mathbf{k_r} - \boldsymbol{k_{ri}})e^{jk_{zi}z} \tag{9}$$

Notice that Eq. (9) can only be used for derivation that follows a vector format, which is typically used in many others[1,2]. Here we will not use it in our derivation since we will focus on separated coordinates $(k_x, k_y, k_z)$.

**Discussion on dispersion parameter $\boldsymbol{\alpha}$**: Meanwhile, ω is the angular frequency of light wave and $\omega_0$ is the central frequency, Δω is the spectral FWHM bandwidth of low coherence light source, $\alpha$ is the dispersion parameters (in unit of mm/Hz), which is proportional to the grating period and also focal length (NA) of tube lens (assuming a unit magnification between grating/DMD and back focal plane of objective). In our system, the diameter of back aperture is $d$ =6.0 mm. When $x$-axis (dispersion axis) overfilling the back aperture $x$-axis, the $\alpha$ parameter is then about:

$$\alpha = \frac{d}{\Delta\omega}\left(\frac{mm}{Hz}\right) \tag{8}$$

which is calculated to be $\alpha = 5.1 \times 10^{-14} mm/Hz$. With broader spectrum, this parameter is smaller than the dispersion parameter demonstrated in a temporal focusing 2P system ($\alpha_{ref} = 1.44 \times 10^{-13} mm/Hz$ from Zhu, Guanghao, et al[3]).

**Discussion on spectrum function**: $S(\omega)$ is the spectrum of low coherence light source, if it is Gaussian distribution:

$$S(\omega) = e^{-\frac{(\omega-\omega_0)^2}{(\Delta\omega)^2}} \tag{9}$$

In fact, in our system, we have a flat spectrum and therefore, $S(\omega)$ is just a step function as:

$$S(\omega) = \begin{cases} 1, |\omega - \omega_0| \leq \Delta\omega/2 \\ 0, \text{otherwise} \end{cases} \tag{10}$$

By simply plug into Renjie et al's derivation[4] of Eq. (A9a), we will conclude to the following equation at plane where camera is conjugate to sample plane $z = 0$:

$$\widetilde{U}_{bs}(k_x, k_y, z_s, \omega) = \frac{\left(\frac{\omega}{c}\right)^2 S(\omega)P(k_x, k_y, \omega)e^{jq(k_x,k_y,\omega)z_R}e^{jk_{zi}(\omega)z_s}}{2q(k_x, k_y, \omega)}$$

$$\cdot\, \tilde{\chi}\big(k_x - k_{xi}(\omega), k_y, -q(k_x, k_y, \omega) - k_{zi}(\omega)\big)$$

where $q(k_x, k_y, \omega) = \sqrt{\left(\frac{n_m\omega}{c}\right)^2 - k_x^2 - k_y^2}$. The pupil function $P(k_x, k_y, \omega)$ is determined as:

$$P(k_x, k_y, \omega)\begin{cases} 1, k_x^2 + k_y^2 < \left(\frac{\omega NA}{c}\right)^2 \\ 0, \text{otherwise} \end{cases} \quad (12)$$

The definitions of $k_{xi}$ and $k_{zi}$ follow the definition from Eq. (5) and (6).

The back-reflected electric fields from reference arm can be written as:

$$\tilde{U}_r(k_x, k_y, z = 0, \omega) = S(\omega)e^{-jk_{xi}\alpha(\omega-\omega_0)}P(k_x, k_y, \omega)\delta(k_x - k_{xi})\delta(k_y) \quad (13)$$

The interference term in the spatial frequency domain is then:

$$\Gamma_{12}(k_x, k_y, z_R, \omega) = \tilde{U}_{bs} \otimes_{\boldsymbol{k_r}} \tilde{U}_r^* \quad (14)$$

$$= \iint \tilde{U}_{bs}(k_x', k_y', z, \omega)\tilde{U}_r(-k_x + k_x', -k_y + k_y', z, \omega)dk_x'dk_y'$$

$$= S^*(\omega)e^{jk_{xi}\alpha(\omega-\omega_0)} \iint \tilde{U}_{bs}(k_x', k_y', z, \omega) \cdot \delta(-k_x + k_x' - k_{xi})\delta(-k_y + k_y')dk_x'dk_y'$$

$$= S^*(\omega)e^{jk_{xi}\alpha(\omega-\omega_0)}\tilde{U}_{bs}(k_x + k_{xi}, k_y)$$

$$= \frac{\left(\frac{\omega}{c}\right)^2 |S(\omega)|^2 P(k_x + k_{xi}, k_y, \omega)e^{jq(k_x+k_{xi},k_y,\omega)z_R}e^{jk_{zi}(\omega)z_s}}{2q(k_x + k_{xi}, k_y, \omega)}$$

$$\cdot\, \tilde{\chi}\left(k_x, k_y, -q(k_x + k_{xi}, k_y, \omega) - k_{zi}(\omega)\right) \quad (15)$$

Notice that $P = |P|^2$ for unit pupil function. And it is interesting that in the end, the lateral profile of illumination pattern actually has nothing to do with the OTF, as it cancels out between two arms ($e^{-jk_{xi}\alpha(\omega-\omega_0)}$ is cancelled out).

For point scatter $\tilde{\chi} = 1$, the above equation reduces to:

$$\Gamma_{12}(k_x, k_y, z_s) = \int_{\omega_0-\Delta\omega/2}^{\omega_0+\Delta\omega/2} \frac{\left(\frac{\omega}{c}\right)^2 |S(\omega)|^2 P(k_x + k_{xi}(\omega), k_y, \omega)e^{j\{q[k_x+k_{xi}(\omega),k_y,\omega]+k_{zi}(\omega)\}z_s}}{2q[k_x + k_{xi}(\omega), k_y, \omega]} d\omega \quad (16)$$

This is our 3D OTF. We can numerically simulate the OTF by using this equation.

We will analyze this OTF from two approaches.

**(I1)** Special case at $\boldsymbol{k_r} = 0$:

Since the bandwidth of low-coherence light source we are using is about $\Delta\omega = 40\ nm$, which is much smaller than $\lambda = 800\ nm$(with 40/800 $\sim$5%), therefore the difference between angular frequency is trivial and then we can assume that $\omega \approx \omega_0$. Also assume a flat spectrum of light source, that's $|S(\omega)| = 1$. Then it will be reduced to:

$$\Gamma_{12}(z_s) = \left(\frac{\omega_0}{c}\right)^2 \int_{-\Delta\omega/2}^{\Delta\omega/2} \frac{P(k_{xi})e^{j2k_{zi}(\delta\omega)z_s}}{2k_{zi}(\delta\omega)} d(\delta\omega) \tag{17}$$

where $\delta\omega = \omega - \omega_0$. The definition of $k_{xi}$ will be:

$$k_{xi} = \frac{\alpha\omega_0}{cf_0} \cdot \delta\omega \tag{18}$$

Since $\alpha = d/\Delta\omega$, it leads to $k_{xi} = \frac{d\omega_0}{\Delta\omega c f_0} \cdot \delta\omega$. And $|\delta\omega| \le \Delta\omega/2$, therefore, $|k_{xi}| \le \frac{d}{2f_0} \cdot \frac{\omega_0}{c}$, which is equivalent to $|k_{xi}| \le k_0 NA$.

And:

$$k_{zi} = \sqrt{\beta_0^2 - k_{xi}^2} \tag{19}$$

with $\beta_0 = \frac{n_m\omega_0}{c} = k_0 n_m$. Therefore,

$$\delta\omega = \left(\beta_0^2 - k_{zi}^2\right) \cdot \frac{cf_0}{\alpha\omega_0} \tag{20}$$

$$d(\delta\omega) = -\frac{cf_0}{\alpha\omega_0} \cdot 2k_{zi}dk_{zi} \tag{21}$$

Let $2k_{zi} = K$. Since $k_{xi} = \sqrt{\beta_0^2 - \left(\frac{K}{2}\right)^2}$, which makes the pupil function $P(k_{xi})$ equivalent to $P(K)$ with $K < \beta_0$. With $k_{xi} < k_0 NA$, then $2k_0\sqrt{n_m^2 - NA^2} < K$. Now it is implied to:

$$\Gamma_{12}(z_R) \propto \int e^{jKz_R} \cdot P(K)dK = \mathcal{F}\{P(K)\}|_K \tag{22}$$

This is simply a FFT of modified pupil function $P(K)$. Here, $\sqrt{n_m^2 - NA^2} < K/2k_0 < n_m$. The Fourier transform of above is equal to FFT of a shifted step function centralized around zero within a range of $2k_0 \cdot [\frac{\sqrt{n_m^2 - NA^2} - n_m}{2}, \frac{n_m - \sqrt{n_m^2 - NA^2}}{2}]$, i.e., $[-T, T]$ with $T = k_0(n_m - \sqrt{n_m^2 - NA^2})$. Therefore, it concludes to:

$$\Gamma(z_s) = 2T \cdot sinc(z_s T) \tag{23}$$

with FWHM $z_{FWHM} = 1.2\pi/T$, therefore,

$$z_{FWHM} = \frac{1.2\pi}{k_0\left(n_m - \sqrt{n_m^2 - NA^2}\right)} \tag{24}$$

It gives axial resolution of $z_{FWHM} \sim 1.06\ \mu m$ with $NA = 1.0$ and wavelength $\lambda_0 = 800\ nm$.

**(I2)** The numerical simulation results are discussed as follows.

(1) 3D OTF: the simulation 3D OTF of speckle system is on the left while our temporal focusing system is on the right. The line profile is the profile along $k_r = 0$.

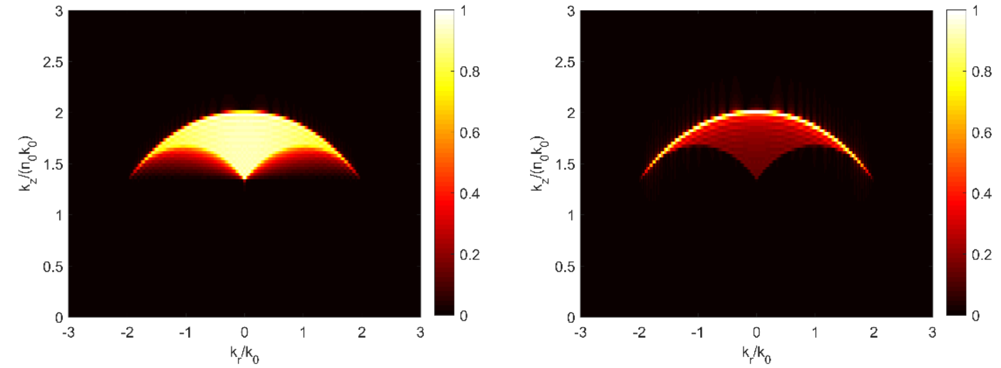

Figure S1: 3D OTF in $(k_z, k_r)$ space

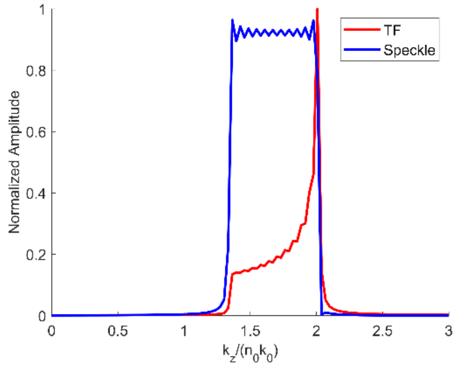

Figure S2: Line profile of OTF$(k_z, k_r = 0)$

(2) PSF along $(x, z)$. The left is speckle system while right is our system.

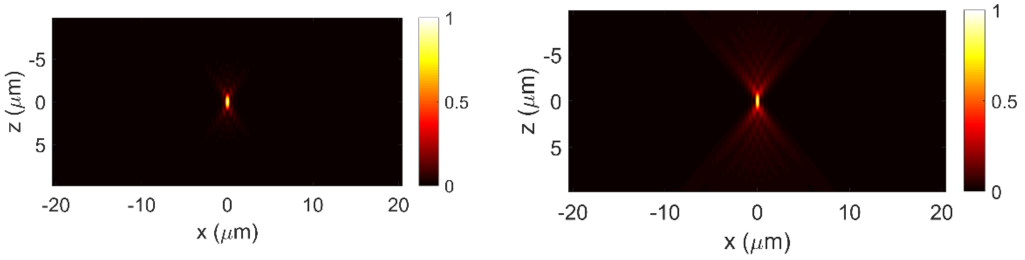

Figure S3: PSF along $(z, x)$ plane

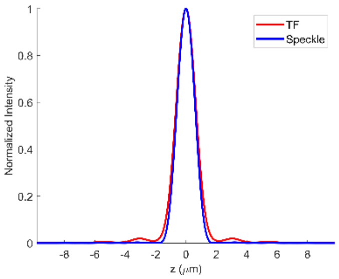

Fig S4: Axial response, both FWHM with Gaussian fitted to be around 1.0 $\mu m$

#### Section II: Pulse width cross-correlation

The temporal focusing enabled gating mechanism resulting from the cross-correlation of different pulses width at different depths. Due to the dispersion, pulse width will be narrowest at the focal plane while broadened as defocused. The interference between sample and reference arms are actually a cross-correlation between the pulse widths at different depths. Therefore, it doesn't matter what is the pulse width. What matters here is how pulse width is broadened relative to focal plane, which is determined by the temporal focusing behavior as multiphoton microscopy application (see back aperture of the objective).

To be noticed, OCT is very different from what we are doing. For OCT imaging system, the pulse width remains constant (if no sample dispersion introduced) and therefore, the axial resolution is simply relied on the temporal coherence length itself. In our system, the pulse widths are depth-dependent, and the function mechanism of axial sectioning is very similar to *autocorrelator used for measuring the ultrashort pulse width*.

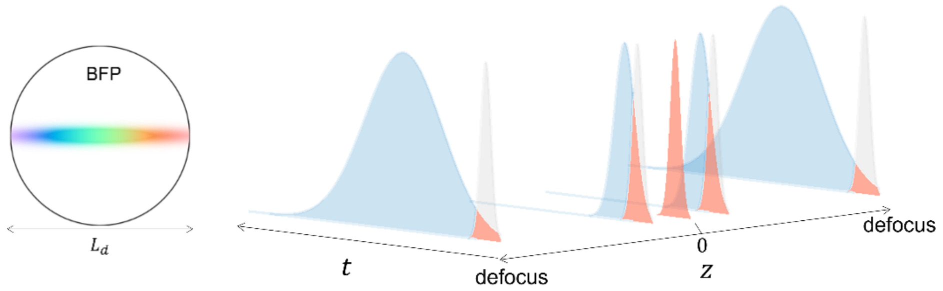

Fig S5. (left) Illumination at back aperture of objectives; (right) pulse width broadened when defocused, the defocused pulse will decorrelated from pulse at focal plane

For pulse width (as pulsed interferometry), the reference arm needs additional term as:

$$e^{-a_0 t^2} \tag{25}$$

as $a_0 = 2ln2/\tau_0^2$, where $\tau_0$ is the pulse width at focal plane.

The sample arm has term:

$$e^{-a(z_s)(t-t_s)^2} \tag{26}$$

where $a(z_s) = 2ln2/\tau^2(z_s)$ is the depth-dependent pulse width in sample arm. The cross-correlation integration will integral over each pulse width duration at each sample plane, therefore, result in a pulsed interferometry. Only consider simple equation, leading to:

$$U(r, z_s) = \int_{-\infty}^{+\infty} U_{bs}(r, z_s) e^{-a(z_s)(t-t_s)^2} \cdot U_R^* e^{-a(z_s|_{z_s=0})t^2} dt \quad (27)$$

This is the cross-correlation between the pulse width, and therefore, giving rise to the axial sectioning performance. This integral indicates the results of cross-correlation between two Gaussian functions. If considering the plane wave illumination with an axially scanning mirror in sample arm while a fixed high-reflective mirror in reference arm, the resulting-in normalized cross-correlation is:

$$U(r, z_s) = |U_{bs}(r)||U_R(r)| \cdot \sqrt{\frac{2\tau(z_s)\tau_0}{\tau^2(z_s) + \tau_0^2}} \cdot e^{-\frac{(2ln2\cdot\tau_s)^2}{\tau^2(z_s)+\tau_0^2}} \quad (28)$$

where $z_s = t_s c/n_m$ with $c$ is light speed while $n_m$ is the refractive index of medium. The full-width half-maximum (FWHM) is then:

$$FWHM_z = 2\sqrt{ln2} \cdot \sqrt{\tau^2(z_s) + \tau_0^2} \quad (29)$$

Apparently, the axial resolution is determined by the decorrelation of pulse widths itself. According to Papagiakoumou Eirini et al[5], the depth-dependent pulse broadening is:

$$\tau(z_s) = \frac{2\sqrt{2ln2}}{\Omega} \sqrt{1 + \frac{z_M}{z_B} \cdot \frac{z_s^2}{z_s^2 + z_M z_R}} \quad (30)$$

with $z_B = 2f_{ob}^2/k_0\alpha^2\Omega^2$, $z_M = 2f_{ob}^2/k_0 s^2$ and $z_R = 2f_{ob}^2/k_0(s^2 + \alpha^2\Omega^2)$ are chirped focusing parts, Rayleigh lengths for the spatial, the spatial and chirped (spatiotemporal) parts of the beam, respectively. $2\sqrt{2ln2}s$ is the FWHM of each monochromatic beam at the back aperture of objective, $k_0$ is the wavevector of the central frequency of the pulse, $\alpha$ is the constant proportional to the grating groove density and the focal length of the collimating lens, $\Omega$ is the FWHM of the frequency spectrum of the pulse and $f_{ob}$ is the objective focal length. $\alpha\Omega$ is the linear spatial chirp at the back aperture, i.e. the extension of the illumination at the objective back aperture in the chirping direction where the wavelength components are dispersed.

Typically, $s \ll \alpha^2\Omega^2$, and $z_R \sim z_B$, and $z_M \gg z_R$, the equation can be:

$$\tau(z_s) \sim \tau_0 \sqrt{1 + \frac{z_S^2}{z_R^2}} \quad (31)$$

$$z_R \sim \frac{2f_{ob}^2}{k_0\alpha^2\Omega^2} \quad (32)$$

The chirp coefficient $\alpha\Omega \sim d$, where $d$ is the diameter of back aperture of the objective. Then,

$$z_R \sim \frac{1}{2k_0 NA^2} \tag{33}$$

Therefore,

$$\tau(z_s) \sim \tau_0 \sqrt{1 + (2k_0 NA^2)^2 \cdot z_s^2} \tag{34}$$

The decorrelation of $\tau(z_s)$ is directly proportional to $2k_0 NA^2$, indicating magnitude of the optical sectioning is similar to most confocal microscope.

### Section III: Discussion of speckle-free coherent imaging

The temporal focusing QPM system can intrinsically achieve speckle-free coherent imaging due to the angular compounding approaches. Since each illumination spot at back-aperture of objectives are independent from each other, each illumination angles are uncorrelated from each other. Each camera exposure will integral all the photons captured from all the angles, where all the angles are independent from each other, and therefore, averaging out.

One interesting observation reported in paper published by Papagiakoumou Eirini et al[6], which described a phenomenon of temporal focusing laser beams with an intrinsic resistance to scattering, namely 'self-healing' properties. This phenomenon is illustrated as following figure. This phenomenon describes two interesting facts: (1) each wavelength component illuminate sample from different illumination angles. At presence of scattering medium, wavelength components that follow propagation path far apart from one another generate uncorrelated speckle patterns. Summing these speckle patterns results in an overall smoothing of the image, therefore be 'speckle-free'; (2) Wavelength components that are adjacent to each other generate highly correlated but only spatially shifted speckles, leading to so-called 'anisotropic smoothing in the direction of the shift'. This effect gives rise to a 'self-healing' property, that's, in my understanding, the distortion or loss of energy due to light scattering at one wavelength will be compensated by the adjacent wavelength component. They have demonstrated that such a temporal focusing beam can penetrate into depth as deep as 200 $\mu m$ inside brain slice.

The speckle-free performance can also be described as following: each illumination angle has a reflected interference signal as:

$$U_i(\boldsymbol{r}, \boldsymbol{r_i}) = U_s(\boldsymbol{r}, \boldsymbol{r_i}) U_r^*(\boldsymbol{r}, \boldsymbol{r_i}) + U_s(\boldsymbol{r}, \boldsymbol{r_i}) U_r^*(\boldsymbol{r}, \boldsymbol{r_i}) \mathrm{e}^{j\varphi(\boldsymbol{r}, \boldsymbol{r_i})} \tag{36}$$

where the first part of right-hand equation represents desired signals while the second part represents speckle noises (cross-talk or interference from different scatters inside the imaging slice due to the spatial coherence of light source). The speckle noises show random phases. For a single illumination angles or any other illumination that results in correlated speckles,

$$U_s(\boldsymbol{r}, \boldsymbol{r_i}) U_r^*(\boldsymbol{r}, \boldsymbol{r_i}) \mathrm{e}^{j\varphi(\boldsymbol{r}, \boldsymbol{r_i})} = U_r^*(\boldsymbol{r}, \boldsymbol{r_i}) \cdot \frac{1}{M} \sum_{k=1}^{M} U_s(r_k, r_i) e^{j\varphi(\boldsymbol{r_k}, \boldsymbol{r_i})} \tag{37}$$

where $k$ indicates scattering signals resulting from other scatters and $M$ is the total number of scatters. Each scattering signals contains contribution from single scatter and multiple scatters, as well as scattering signals from other pixels. Due to the cross-correlation between those scattering signals, it will contribute to the speckle noises (in a form of summation over the amplitude field).

However, with multiple independent illumination angles, the speckle patterns are averaged over intensity field,

$$U(\boldsymbol{r}) = \frac{1}{N}\sum_{i=1}^{N}\left|U_s(\boldsymbol{r},\boldsymbol{r_i})U_r^*(\boldsymbol{r},\boldsymbol{r_i})\mathrm{e}^{j\varphi(r,r_i)}\right|^2 \tag{38}$$

which will ultimately smooth out the speckle noise and therefore the speckle noise will be greatly reduced.

#### Section IV: Trade-off between field of view and spectral bandwidth

In TF-QPM, the plane of grating is conjugated to the focal plane of objective where the achievable FOV is then determined by the total magnification factor between the two planes. The smaller magnification it is, the larger FOV it can achieve. In contrast, to make full of the NA of objective, it is necessary to overfill the back aperture of the objective. In principle, it should satisfy the following condition[7]:

$$\frac{f \cdot \Delta\lambda \cdot k_g}{D} \sim 1 \tag{39}$$

where $f$ is the effective focal length of lens system between the grating and back aperture plane of the objective, $\Delta\lambda$ is the spectral bandwidth of light source, $k_g$ (lines/mm) is the grating period and $D$ is the back aperture diameter of the objective.

On the other hand, the achievable imaging FOV is proportional to the magnification factor between the systems, which is defined as:

$$M = f/f_{TL} \cdot M_{obj} \tag{40}$$

where $f_{TL}$ is the focal length of the designed tube length of objective which is 180 mm for Olympus objective and $M_{obj}$ is the designed magnification factor of the objective. In our case, we use Olympus 60x objective with $D\ = 6\,mm$ and $M_{obj} = 60$. The maximum achievable FOV is then defined as ($D_{grating}$ is the smallest dimension of the grating, assuming beam is overfilling the grating):

$$2r = D_{grating}/M \propto 1/f \tag{41}$$

With a fixed $D$ and $k_g\ = 1200$ lines/mm, we also have

$$\Delta\lambda \propto 1/f \tag{42}$$

Therefore, with a smaller focal length (magnification) of lens system after grating, we can achieve a larger imaging FOV but would require larger spectral bandwidth to achieve axial sectioning making full use of NA of the objective. The simulated $\Delta\lambda\ vs. M$ as well as $2r\ vs. M$ is shown in Fig. S6.

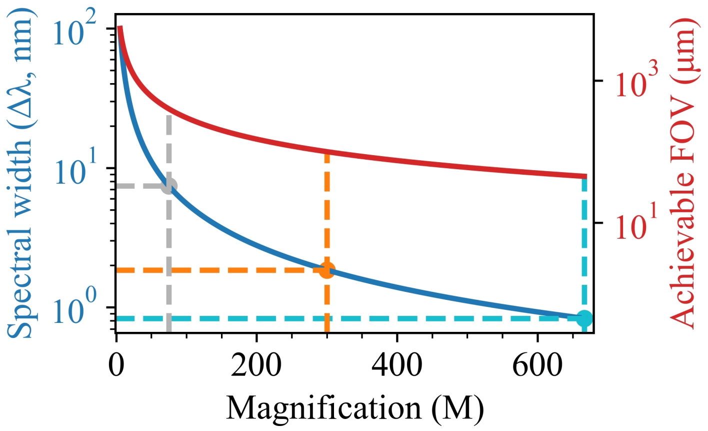

Fig. S6: simulation the tradeoff between spectral bandwidth and achievable FOV. Both y-axes are in log scale.

To achieve imaging FOV of 400 $\mu m$, 100 $\mu m$ and 45 $\mu m$, it requires magnification of 75, 300, and 667, with spectral bandwidth of 7.4 nm, 1.8 nm, and 0.8 nm, respectively. Though the above simulation assumes an infinite large aperture of the lens system, the geometric size of the lenses will ultimately impose additional limitation to the available maximum FOV. From planes of grating to focal plane of objective, the aperture of lens system should be able to collect both the spatial beam width and the angular spectral fan simultaneously. Otherwise, vignetting by the lens system will clip the off-axis spectral components. The angular dispersion of the laser beam is approximately (at diffraction order of *m*):

$$\Delta\theta_{spectral} = \frac{k_g m}{cos\theta}\Delta\lambda_{laser} \tag{43}$$

The aperture (radius) of the lens system should be able to collect the beam as large as:

$$R_{lens} \geq \frac{D_{grating}}{2} + f \cdot \frac{\Delta\theta_{spectral}}{2} \tag{44}$$

where $D_{grating}$ is the smallest dimension of the grating and we assume that the light overfills the grating. If the geometric constrain set the maximum achievable FOV, then the maximum achievable FOV in object space would then be:

$$2r \sim \frac{D_{grating}}{M} \leq \frac{2R_{lens}}{M} - f \cdot \frac{\Delta\theta_{spectral}}{2} \tag{45}$$

A transform-limited Ti:Sapphire laser usually provides ~nm order of bandwidth while a picosecond laser with pm to sub-nm order of the bandwidth. Therefore, to achieve maximum FOV in TF-QPM, the options of light source should be taken into consideration.

#### Section V: Comparison different PIV techniques

| Technique | 3D? | Frame rate | Axial resolution | Lateral resolution | FOV | Tracers* | Penetration depth** |
|---|---|---|---|---|---|---|---|
| Planar PIV[8] | 2D | 10-50 kHz | 0.5-2 mm | 0.1 – 1 mm | 10 x 10 $mm^2$ | S | shallow |
| Micro-PIV (epi-fluorescence[9]) | 2D | 1-5 kHz | 10-100 um | 1-10 μm | 0.2-2 $mm^2$ | F | shallow |

| | | | | | | | |
|---|---|---|---|---|---|---|---|
| Confocal scanning PIV[10] | 2D-3D | 2 kHz | 1.88 µm | 9.1 µm | 288 x 171 $\mu m^2$ | F | **deep** |
| Tomographic PIV[11] | 3D | 1-5 kHz | 20-200 µm | 0.5 – 2 mm | 50-100 $mm^3$ | S | shallow |
| Digital holographic PIV[12] | 3D | 10s Hz | 1-10 µm | 0.1 – 1 mm | Sub-mm | S | shallow |
| Ultrasound PIV[13] | 3D | 10 – 1000 Hz | 50 – 300 µm | 100 – 500 µm | mm | S | **deep** |
| TF-QPM (our approach) | 2D-3D | 3 kHz in 2D (camera limited) | < 1 µm | < 0.5 µm | 100 x 80 $\mu m^2$ | S | **deep** |

*S: scattering particle (for ultrasound, it is micro-size gas bubble); F: fluorescence particles

**Planar PIV, micro-PIV, tomographic and digital holographic PIV are based on transmission geometry, making it unsuitable for thick opaque sample where certain penetration depth is required. Confocal PIV and our TF-QPM are based on reflection mode with depth selectivity, enabling the potential for in vivo imaging through thick opaque sample, like tissues.
